# MSA Transformer

**DOI:** 10.1101/2021.02.12.430858

**Authors:** Roshan Rao, Jason Liu, Robert Verkuil, Joshua Meier, John F. Canny, Pieter Abbeel, Tom Sercu, Alexander Rives

**Affiliations:** UC Berkeley; Work performed during internship at FAIR; Facebook AI Research; New York University

## Abstract

Unsupervised protein language models trained across millions of diverse sequences learn structure and function of proteins. Protein language models studied to date have been trained to perform inference from individual sequences. The longstanding approach in computational biology has been to make inferences from a family of evolutionarily related sequences by fitting a model to each family independently. In this work we combine the two paradigms. We introduce a protein language model which takes as input a set of sequences in the form of a multiple sequence alignment. The model interleaves row and column attention across the input sequences and is trained with a variant of the masked language modeling objective across many protein families. The performance of the model surpasses current state-of-the-art unsupervised structure learning methods by a wide margin, with far greater parameter efficiency than prior state-of-the-art protein language models.

## 1. Introduction

Unsupervised models learn protein structure from patterns in sequences. Sequence variation within a protein family conveys information about the structure of the protein (Yanofsky et al., 1964; Altschuh et al., 1988; Göbel et al., 1994). Since evolution is not free to choose the identity of amino acids independently at sites that are in contact in the folded three-dimensional structure, patterns are imprinted onto the sequences selected by evolution. Constraints on the structure of a protein can be inferred from patterns in related sequences. The predominant unsupervised approach is to fit a Markov Random Field in the form of a Potts Model to a family of aligned sequences to extract a coevolutionary signal (Lapedes et al., 1999; Thomas et al., 2008; Weigt et al., 2009).

A new line of work explores unsupervised protein language models (Alley et al., 2019; Rives et al., 2020; Heinzinger et al., 2019; Rao et al., 2019). This approach fits large neural networks with shared parameters across millions of diverse sequences, rather than fitting a model separately to each family of sequences. At inference time, a single forward pass of an end-to-end model replaces the multistage pipeline, involving sequence search, alignment, and model fitting steps, standard in bioinformatics. Recently, promising results have shown that protein language models learn secondary structure, long-range contacts, and function via the unsupervised objective (Rives et al., 2020), making them an alternative to the classical pipeline. While small and recurrent models fall well short of state-of-the-art (Rao et al., 2019), the internal representations of very large transformer models are competitive with Potts models for unsupervised structure learning (Rives et al., 2020; Rao et al., 2021).

Potts models have an important advantage over protein language models during inference. The input to the Potts model is a set of sequences. Inference is performed by fitting a model that directly extracts the covariation signal from the input. Current protein language models take a single sequence as input for inference. Information about evolutionary variation must be stored in the parameters of the model during training. As a result, protein language models require many parameters to represent the data distribution well.

In this work, we unify the two paradigms within a protein language model that takes sets of aligned sequences as input, but shares parameters across many diverse sequence families. Like prior protein language models operating on individual sequences, the approach benefits from learning from common patterns across protein families, allowing information to be generalized and transferred between them. By taking sets of sequences as input, the model gains the ability to extract information during inference, which improves the parameter efficiency.

We introduce the MSA Transformer, a model operating on sets of aligned sequences. The input to the model is a multiple sequence alignment. The architecture interleaves attention across the rows and columns of the alignment as in axial attention (Ho et al., 2019; Huang et al., 2019). We propose a variant of axial attention which shares a single attention map across the rows. The model is trained using the masked language modeling objective. Self supervision is performed by training the model to reconstruct a corrupted MSA.

We train an MSA Transformer model with 100M parameters on a large dataset (4.3 TB) of 26 million MSAs, with an average of 1192 sequences per MSA. The resulting model surpasses current state-of-the-art unsupervised structure learning methods by a wide margin, outperforming Potts models and protein language models with 650M parameters. The model improves over state-of-the-art unsupervised contact prediction methods across all multiple sequence alignment depths, with an especially significant advantage for MSAs with lower depth. Information about the contact pattern emerges directly in the tied row attention maps. Evaluated in a supervised contact prediction pipeline, features captured by the MSA Transformer outperform trRosetta (Yang et al., 2019) on the CASP13 and CAMEO test sets. We find that high precision contact predictions can be extracted from small sets of diverse sequences, with good results from as few as 8-16 sequences. We investigate how the model performs inference by independently destroying the covariation or sequence patterns in the input, finding that the model uses both signals to make predictions.

## 2. Related Work

### Unsupervised Contact Prediction

The standard approach to unsupervised protein structure prediction is to identify pairwise statistical dependencies between the columns of an MSA, which are modeled as a Potts model Markov Random Field (MRF). Since exact inference is computationally intractable, a variety of methods have been proposed to efficiently fit the MRF, including mean-field inference (Morcos et al., 2011), sparse-inverse covariance estimation (Jones et al., 2012), and the current state-of-the-art, pseudolikelihood maximization (Balakrishnan et al., 2011; Ekeberg et al., 2013; Seemayer et al., 2014). In this work we use Potts models fit with psuedolikelihood maximization as a baseline, and refer to features generated from Potts models as “co-evolutionary features.” Making a connection with the attention mechanism we study here, Bhattacharya et al. (2020) show that a single layer of self-attention can perform essentially the same computation as a Potts model.

### Deep Models of MSAs

Several groups have proposed to replace the shallow MRF with a deep neural network. Riesselman et al. (2018) train deep variational autoencoders on MSAs to predict function. Riesselman et al. (2019) train autoregressive models on MSAs, but discard the alignment, showing that function can be learned from unaligned sequences. In contrast to our approach which is trained on many MSAs, these existing models are trained on a single set of related sequences and do not provide a direct method of extracting protein contacts.

### Supervised Structure Prediction

Supervised structure prediction using deep neural networks has driven ground-breaking progress on the protein structure prediction problem (Senior et al., 2019; Jumper et al., 2020). Initial models used coevolutionary features (Wang et al., 2017; Liu et al., 2018; Yang et al., 2019; Senior et al., 2019; Adhikari & Elofsson, 2020). Recently MSAs have been proposed as input to supervised structure prediction methods. Mirabello & Wallner (2019) and Kandathil et al. (2020) study models that take MSAs as input directly, respectively using 2D convolutions or GRUs to process the input. More recently, AlphaFold2 (Jumper et al., 2020) uses attention to process MSAs in an end-to-end model supervised with structures.

The central difference in our work is to model a collection of MSAs using *unsupervised learning*. This results in a model that contains features potentially useful for a range of downstream tasks. We use the emergence of structure in the internal representations of the model to measure the ability of the model to capture biology from sequences. This is a fundamentally distinct problem setting from supervised structure prediction. The MSA Transformer is trained in a purely unsupervised manner and learns contacts without being trained on protein structures.

Large protein sequence databases contain billions of sequences and are undergoing exponential growth. Unsupervised methods can directly use these datasets for learning, while supervised methods are limited to supervision from the hundreds of thousands of crystallized structures. Unsupervised methods can learn from regions of sequence space not covered by structural knowledge.

### Protein Language Models

Protein language modeling has emerged as a promising approach for unsupervised learning of protein sequences. **?** combined unsupervised sequence pre-training with structural supervision to produce sequence embeddings. Alley et al. (2019) and Heinzinger et al. (2019) showed that LSTM language models capture some biological properties. Simultaneously, Rives et al. (2020) proposed to model protein sequences with self-attention, showing that transformer protein language models capture accurate information of structure and function in their representations. Rao et al. (2019) evaluated a variety of protein language models across a panel of benchmarks concluding that small LSTMs and transformers fall well short of features from the bioinformatics pipeline.

A combination of model scale and architecture improvements has been instrumental to recent successes in protein language modeling. Elnaggar et al. (2020) study a variety of transformer variants. Rives et al. (2020) show that large transformer models produce state-of-the-art features across a variety of tasks. Notably, the internal representations of transformer protein language models are found to directly represent contacts. **?** find that specific attention heads of pre-trained transformers correlate directly with protein contacts. Rao et al. (2021) combine multiple attention heads to predict contacts more accurately than Potts models, despite using just a single sequence for inference.

Alternatives to the masked language modeling objective have also been explored, such as conditional generation (Madani et al., 2020) and contrastive loss functions (Lu et al., 2020). Most relevant to our work, Sturmfels et al. (2020) and Sercu et al. (2020) study alternative learning objectives using sets of sequences for supervision. Sturmfels et al. (2020) extended the unsupervised language modeling to predict the position specific scoring matrix (PSSM) profile. Sercu et al. (2020) used amortized optimization to simultaneously predict profiles and pairwise couplings. In natural language processing, recent work (Lewis et al., 2020; Gu et al., 2018) has explored models using multiple sequences. However, previous work on protein language models has not considered inference *directly* from sets of sequences.

## 3. Methods

Transformers are powerful sequence models capable of passing information from any position to any other position (Vaswani et al., 2017). However, they are not trivially applied to a set of aligned sequences. Naively concatenating *M* sequences of length *L* in an MSA would allow attention across all sequences, but the (*ML*)^2^ self-attention maps would be prohibitively memory-intensive. The main contribution of this paper is to extend transformer pre-training to operate on an MSA, while respecting its structure as an *M* × *L* character matrix.

We describe the input MSA as a matrix ***x*** ∈ ℝ^*M×L*^, where rows correspond to sequences in the MSA, columns are positions in the aligned sequence, and entries ***x***_*mi*_ take integer values^1^ encoding the amino acid identity of sequence *m* at position *i*. After embedding the input, each layer has a ℝ^*M×L×d*^ state as input and output. For the core of the transformer, we adapt the axial attention approach from Ho et al. (2019), Huang et al. (2019), and Child et al. (2019). This approach alternates attention over rows and columns of the 2D state (see Fig. 1). This sparsity pattern in the attention over the MSA brings the attention cost to *O*(*LM*^2^) for the column attention, and *O*(*ML*^2^) for the row attention.

**Figure 1.**
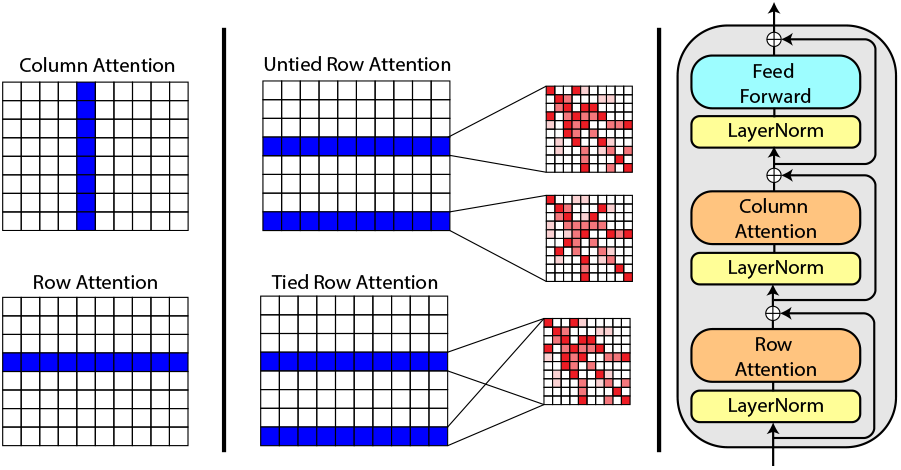
**Left:** Sparsity structure of the attention. By constraining attention to operate over rows and columns, computational cost is reduced from *O*(*M*^2^*L*^2^) to *O*(*LM*^2^) + *O*(*ML*^2^) where *M* is the number of rows and *L* the number of columns in the MSA. **Middle:** Untied row attention uses different attention maps for each sequence in the MSA. Tied row attention uses a single attention map for all sequences in the MSA, thereby constraining the contact structure. Ablation studies consider the use of both tied and untied attention. The final model uses tied attention. **Right:** A single MSA Transformer block. The depicted architecture is from the final model, some ablations alter the ordering of row and column attention.

### Feedforward Layers

We deviate from Ho et al. (2019) in the interleaving of the feedforward layers. Rather than applying a feedforward layer after each row or column attention, we apply row and column attention followed by a single feedforward layer (see Fig. 1). This choice follows more closely the transformer decoder architecture (Vaswani et al., 2017).

### Position Embedding

The standard transformer position embedding is a 1D signal added to each position in the sequence. Either fixed sinusoidal (Vaswani et al., 2017) or learned (Devlin et al., 2019) position embeddings are most commonly used. Rives et al. (2020) found that learned position embeddings generally resulted in better downstream performance for protein language models.

An MSA is a 2D input so we must consider two types of position embeddings. For all models trained, we provide a 1D *sequence* position embedding, which is added independently to each row of the MSA. This allows the model to distinguish different aligned positions. For one model, we also add a position embedding independently to each column of the MSA, which allows the model to distinguish different sequences (without this, the model treats the input sequences as an unordered set). We also ensure that the first position in the sequence is always the reference so that it can always be uniquely identified through the position embedding. We find that incorporating the column position embedding increases performance slightly and so choose to use it in the final model (see Appendix A.3.6 for further discussion).

### Tied Row Attention

The standard implementation of axial attention allows for independent attention maps for each row and column of the input. However, in an MSA each sequence should have a similar structure; indeed, direct-coupling analysis exploits this fact to learn contact information. To leverage this shared structure we hypothesize it would be beneficial to tie the row attention maps between the sequences in the MSA. As an additional benefit, tied attention reduces the memory footprint of the row attentions from *O*(*ML*^2^) to *O*(*L*^2^).

Let *M* be the number of rows, *d* be the hidden dimension and *Q*_*m*_, *K*_*m*_ be the matrix of queries and keys for the *m*-th row of input. We define tied row attention (before softmax is applied) to be:

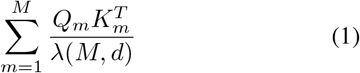

The denominator *λ*(*M, d*) would be the normalization constant 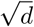 in standard scaled-dot product attention. In tied row attention, we explore two normalization functions to prevent attention weights linearly scaling with the number of input sequences: 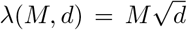 (mean normalization) and 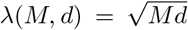 (square-root normalization). Our final model uses square-root normalization.

### Pre-training Objective

We adapt the masked language modeling objective (Devlin et al., 2019) to the MSA setting.

The loss for an MSA ***x***, and masked MSA 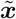 is as follows:

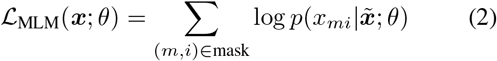

The probabilities are the output of the MSA transformer, softmax normalized over the amino acid vocabulary independently per position *i* in each sequence *m*. We consider masking tokens uniformly at random over the MSA or masking entire columns of the MSA. Masking tokens uniformly at random results in best performance (Table. A.2). Note that the masked token can be predicted not only from context amino acids at different positions but also from related sequences at the same position.

### Pre-training Dataset

Models are trained on a dataset of 26 million MSAs. An MSA is generated for each UniRef50 (**?**) sequence by searching UniClust30 (Mirdita et al., 2017) with HHblits (Steinegger et al., 2019). The average depth of the MSAs is 1192. See Fig. A.2 for MSA depth distribution.

### Models and Training

We train 100M parameters model with 12 layers, 768 embedding size, and 12 attention heads, using a batch size of 512 MSAs, learning rate 10^−4^, no weight decay, and an inverse square root learning rate schedule with 16000 warmup steps. All models are trained on 32 V100 GPUs for 100k updates. The four models with best contact precision are then further trained to 150k updates. Finally, the best model at 150k updates is trained to 450k updates. Unless otherwise specified, all downstream experiments use this model.

Despite the use of axial attention and tied attention to lower the memory requirements, large MSAs still do not easily fit in memory at training time. The baseline model fits a maximum of *N* = 2^14^ tokens on a 32 GB V100 GPU at training time. To work around this limitation we subsample the input MSAs to reduce the number of sequences and limit the maximum sequence length to 1024.

### MSA Subsampling During Inference

At inference time, memory is a much smaller concern. Nevertheless we do not provide the full MSA to the model as it would be computationally expensive and the model’s performance can decrease when the input is much larger than that used during training. Instead, we explore four strategies for subsampling the sequences provided to the model.

- **Random:** This strategy parallels the one used at training time, and selects random sequences from the MSA (ensuring that the reference sequence is always included).
- **Diversity Maximizing:** This is a greedy strategy which starts from the reference and adds the sequence with highest average hamming distance to current set of sequences.
- **Diversity Minimizing:** This strategy is equivalent to the Diversity Maximizing strategy, but adds the sequence with lowest average hamming distance. It is used to explore the effects of diversity on model performance.
- **HHFilter:** This strategy applies hhfilter (Steinegger et al., 2019) with the `-diff M` parameter, which returns *M* or more sequences that maximize diversity (the result is usually close to *M*). If more than *M* sequences are returned we apply the Diversity Maximizing strategy on top of the output.

## 4. Results

We study the MSA Transformer in a panel of structure prediction tasks, evaluating unsupervised contact prediction from the attentions of the model, and performance of features in supervised contact and secondary structure prediction pipelines.

To calibrate the difficulty of the masked language modeling task for MSAs, we compare against two simple prediction strategies using the information in the MSA: (i) column frequency baseline, and (ii) nearest sequence baseline. These baselines implement the intuition that a simple model could use the column frequencies to make a prediction at the masked positions, or copy the identity of the missing character from the most similar sequence in the input. Table. A.1 reports masked language modeling performance. The MSA Transformer model (denoising accuracy of 64.0) significantly outperforms the PSSM (accuracy 41.4) and nearest-neighbor (accuracy 46.7) baselines.

### 4.1. Unsupervised Contact Prediction

Rao et al. (2021) showed that transformer protein language models learned to capture information about protein structure in their attention maps using little to no supervision. This is done by training a small logistic regression (one parameter per attention head) on a limited number of protein structures to predict the probability of a contact between residues *i* and *j* based on the attentions between the residues for all attention heads. The logistic regression weights are shared for all pairs of positions (see Appendix A.1 for more details).

We use the same validation methodology. A logistic regression with 144 parameters is fit on 20 training structures from the trRosetta dataset (Yang et al., 2019). This is then used to predict the probability of protein contacts on another 14842 structures from the trRosetta dataset (training structures are excluded). At inference time, we use hhfilter to subsample 256 sequences.

We compare to two state-of-the-art transformer protein language models: ESM-1b (Rives et al., 2020) with 650M parameters and ProTrans-T5 (Elnaggar et al., 2020) with 3B parameters. For the single-sequence protein language models we use the sequence belonging to the structure as input. We also compare against Potts models using the APC-corrected (Dunn et al., 2008) Frobenius norm of the coupling matrix computed on the MSA (Kamisetty et al., 2013).

Table 1 compares unsupervised contact prediction performance of the models. The MSA Transformer significantly outperforms all baselines, increasing top-L long-range contact precision by a full 15 points over the previous state-of-the-art. Table 2 shows results on harder test sets CAMEO hard targets (Haas et al., 2018) and CASP13-FM (Shrestha et al., 2019). The CASP13-FM test set consists of 31 free modeling domains (from 25 targets); the CAMEO hard targets are a set of 131 domains (out of which we evaluate on the 129 that fit within the 1024 character maximum context length of the model). On the CASP13-FM test set, *unsupevised* contact prediction with the MSA Transformer (43.4 top-L long-range precision) is competitive with the trRosetta base model (45.7 top-L long-range precision), a fully *supervised* structure prediction model.

**Table 1.** Average long-range precision for MSA and single-sequence models on the unsupervised contact prediction task.

| Model | L | L/2 | L/5 |
| --- | --- | --- | --- |
| Potts | 39.3 | 52.2 | 62.8 |
| TAPE | 11.2 | 14.0 | 17.9 |
| ProTrans-T5 | 35.6 | 46.1 | 57.8 |
| ESM-1b | 41.1 | 53.3 | 66.1 |
| MSA Transformer | **57.4** | **71.7** | **82.1** |

**Table 2.** Unsupervised contact prediction on CASP13 and CAMEO (long-range precision). Note the large improvement of MSA Transformer over classical Potts models and ESM-1b.

| Model | CASP13-FM |  | CAMEO |  |
| --- | --- | --- | --- | --- |
|  | L | L/5 | L | L/5 |
| Potts | 16.9 | 31.5 | 24.0 | 42.8 |
| ProTrans-T5 | 16.5 | 27.0 | 25.9 | 43.4 |
| ESM-1b | 17.0 | 30.4 | 30.9 | 52.7 |
| MSA Transformer | **44.8** | **72.5** | **43.5** | **66.8** |

Fig. 2 shows the top-L long-range precision distribution across all structures, comparing the MSA Transformer with Potts models and ESM-1b. The MSA Transformer matches or exceeds Potts models on 98.5% of inputs and matches or exceeds ESM-1b on 91.0% of inputs. Fig. 2 also shows unsupervised contact performance as a function of MSA depth. The model outperforms ESM-1b and Potts models across all MSA depths and has a significant advantage for lower depth MSAs. We find no statistically significant correlation between sequence length and contact precision.

**Figure 2.**
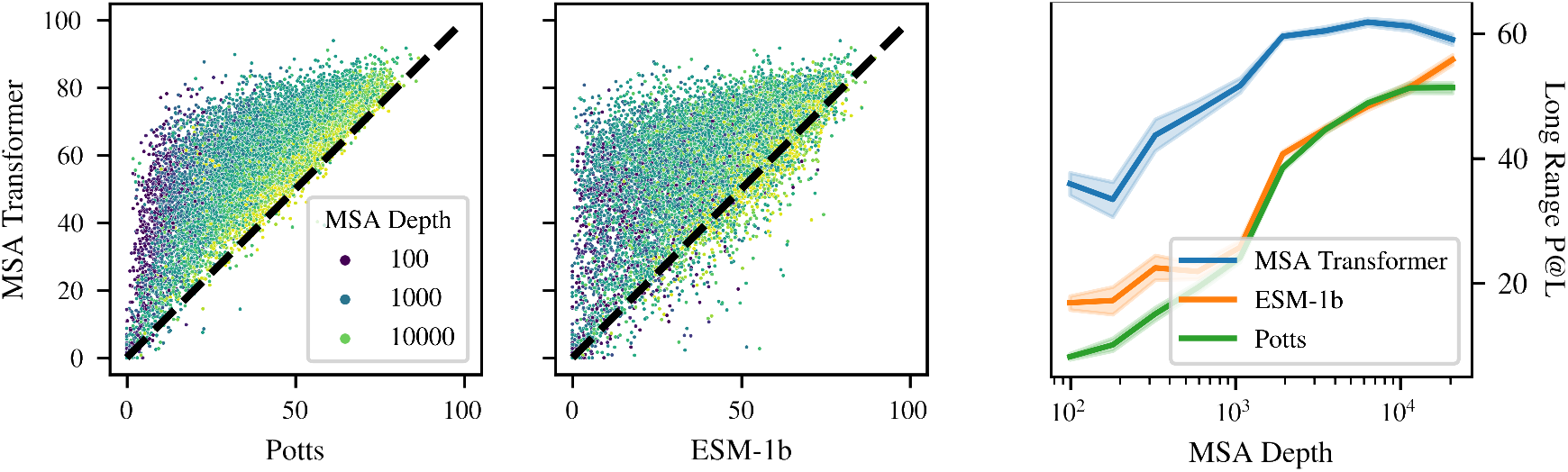
**Left:** Top-L long-range contact precision (higher is better). Comparison of MSA Transformer with Potts model (left scatter plot), and ESM-1b (right scatter plot). Each point represents a single protein (14,842 total) and is colored by the depth of the full MSA for the sequence. The Potts model is given the full MSA as input; ESM-1b is given only the reference sequence; and the MSA Transformer is given an MSA subsampled with hhfilter to a maximum of 256 sequences. The MSA Transformer outperforms both models for the vast majority of sequences. **Right:** Long-range contact precision performance as a function of MSA depth. Sequences are binned by MSA depth into 10 bins; average performance in each bin along with 95% confidence interval is shown. The minimum MSA depth in the trRosetta dataset is 100 sequences. While model performance generally increases with MSA depth, the MSA Transformer performs very well on sequences with low-depth MSAs, rivaling Potts model performance on MSAs 10x larger.

### 4.2. Supervised Contact Prediction

Used independently, features from current state-of-the-art protein language models fall short of co-evolutionary features from Potts models on supervised contact prediction tasks (Rives et al., 2020).

We evaluate the MSA Transformer as a component of a supervised structure prediction pipeline. Following Rives et al. (2020), we train a deep residual network with 32 pre-activation blocks, each with a filter size of 64, using learning rate 0.001. The network is supervised with binned pairwise distance distributions (distograms) using the trRosetta training set (Yang et al., 2019) of 15051 MSAs and structures.

We evaluate two different ways of extracting features from the model. In the first, we use the outer concatenation of the output embeddings of the query sequence. In the second, we combine the outer concatenation with the symmetrized row self-attention maps. For comparison, we train the same residual network over co-evolutionary features from Potts models (Seemayer et al., 2014). Additionally we compare to features from state-of-the-art protein language models ESM-1b and ProTrans-T5 using the outer concatenation of the sequence embeddings. Dropout of 0.1 is used for all language model-based contact predictors. We also compare to trRosetta (Yang et al., 2019), a state-of-the-art supervised structure prediction method prior to AlphaFold2 (Jumper et al., 2020).

The MSA Transformer produces a substantial improvement over co-evolutionary features for supervised contact prediction. Table 3 shows a comparison between the models on the CASP13-FM and CAMEO test sets. The best MSA Transformer model, using the combination of attention maps with features from the final hidden layer, outperforms all other models including the trRosetta baseline model (which uses 36 residual blocks) and the trRosetta full model (which uses 61 residual blocks, data augmentation via MSA subsampling, and predicts inter-residue orientations). Model ensembling over all 5 released models is used in the evaluation of the trRosetta models. Table. A.4 gives additional comparisons with LSTM and transformer protein language models available in the literature.

**Table 3.** Supervised contact prediction on CASP13 and CAMEO (long-range precision). *Uses outer-concatenation of the query sequence representation as features. ^†^Additionally uses the row attention maps as features.

| Model | CASP13-FM |  | CAMEO |  |
| --- | --- | --- | --- | --- |
|  | L | L/5 | L | L/5 |
| trRosetta<sub>base</sub> | 45.7 | 69.6 | 50.9 | 75.5 |
| trRosetta<sub>full</sub> | 51.8 | 80.1 | 53.2 | 77.5 |
| Co-evolutionary | 40.1 | 65.2 | 47.3 | 72.1 |
| ProTrans-T5 | 25.0 | 41.4 | 40.8 | 63.3 |
| ESM-1b | 28.2 | 50.2 | 44.4 | 68.4 |
| MSA Transformer* | 54.5 | **80.2** | 53.6 | 78.0 |
| MSA Transformer† | **54.6** | 77.5 | **55.8** | **79.1** |

### 4.3. Secondary Structure Prediction

To further evaluate the quality of representations generated by the MSA Transformer, we train a state-of-the-art down-stream head based on the Netsurf architecture (Klausen et al., 2019). The downstream model is trained to predict 8-class secondary structure from the pretrained representations. We evaluate models on the CB513 test set (Cuff & Barton, 1999). The models are trained on the Netsurf training dataset. Representations from the MSA Transformer (72.9%) surpass the performance of HMM profiles (71.2%) and ESM-1b embeddings (71.6%) (Table 4).

**Table 4.** CB513 8-class secondary structure prediction accuracy.

| Model | CB513 |
| --- | --- |
| Netsurf | 72.1 |
| HMM Profile | 71.2 ± 0.1 |
| ProTrans-T5 | 71.4 ± 0.3 |
| ESM-1b | 71.6 ± 0.1 |
| MSA Transformer | **73.4 ± 0.3** |

### 4.4. Ablation Study

We perform an ablation study over seven model hyperparameters, using unsupervised contact prediction on the validation set for evaluation. For each combination of hyperparameters, a model is pre-trained with the masked language modeling objective for 100k updates. Training curves for the models are shown in Fig. A.3 and Top-L long-range precision is reported in Table. A.2.

The ablation studies show the use of tied attention plays a critical role in model performance. After 100k updates, a model trained with square-root normalized tied attention outperforms untied attention by more than 17 points and outperforms mean normalized tied-attention by more than 6 points on long-range contact precision.

Parameter count also affects contact precision. A model with half the embedding size (384) and only 30M parameters reaches a long-range precision of 52.8 after 100k updates, 3.5 points lower than the base model, yet 11.7 points higher than the state-of-the-art 650M parameter single-seequence model. See Appendix A.3 for further discussion.

## 5. Model Analysis

We examine how the model uses its input MSA in experiments to understand the role of sequence diversity, attention patterns, and covariation in the MSA.

### 5.1. Effect of MSA diversity

The diversity of the input sequences strongly influences inference of structure. We explore three inference time strategies to control the diversity of the input sequence sets: (i) diversity maximizing, (ii) diversity minimizing, and (iii) random selection (see Section 3). Fig. 4 shows average performance across the test set for each selection strategy as the number of sequences used for input increases. Two approaches to maximize diversity, greedy hamming distance maximization and hhfilter, perform equivalently. Both strategies surpass ESM-1b performance with just 16 input sequences. In comparison, the diversity minimizing strategy, hamming distance minimization, performs poorly, requiring 256 sequences to surpass ESM-1b. Random selection performs well, although it falls behind the diversity maximizing strategies. The qualitative effects of MSA diversity are illustrated in Fig. 3, where the addition of just one high-diversity sequence outperforms the addition of 31 low-diversity sequences.

**Figure 3.**
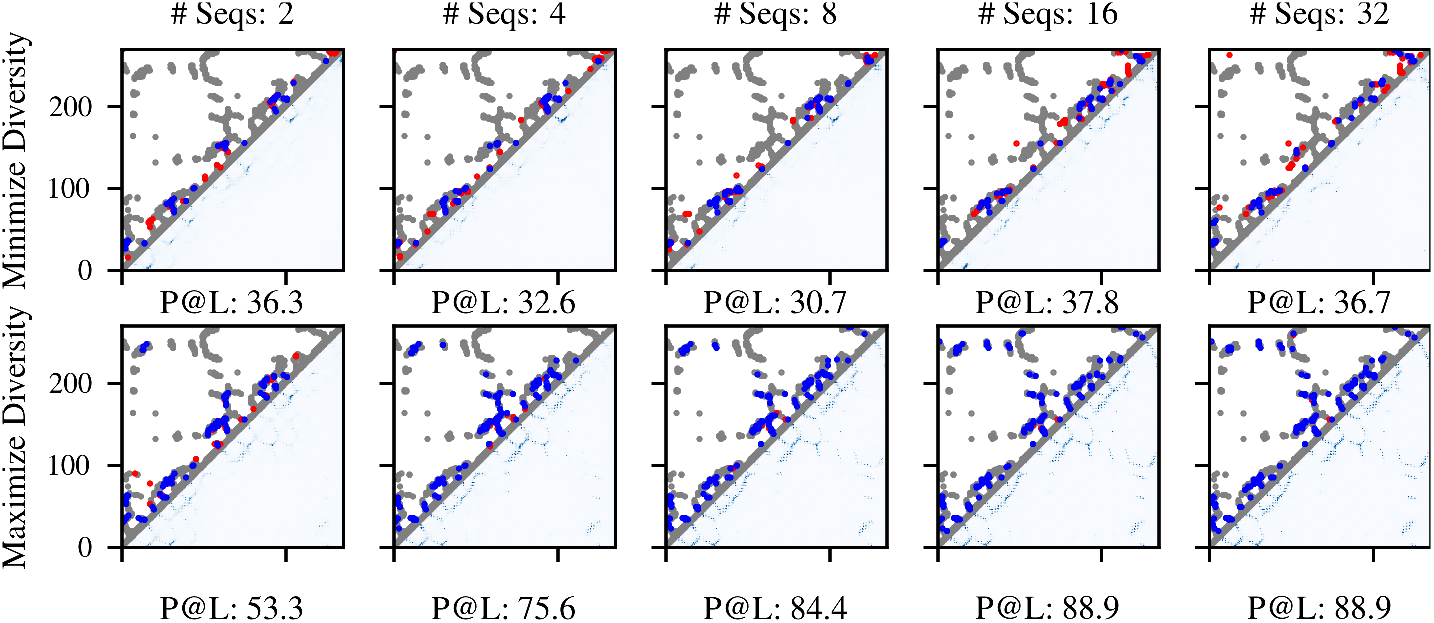
Contact prediction from a small set of input sequences. Predictions are compared under diversity minimizing and diversity maximizing sequence selection strategies. Visualized for 4zjp chain A. Raw contact probabilities are shown below the diagonal, top L contacts are shown above the diagonal. (blue: true positive, red: false positive, grey: ground-truth contacts). Top-L long-range contact precision below each plot. Contact precision improves with more sequences under both selection strategies. Maximizing the diversity enables identification of long-range contacts from a small set of sequences.

**Figure 4.**
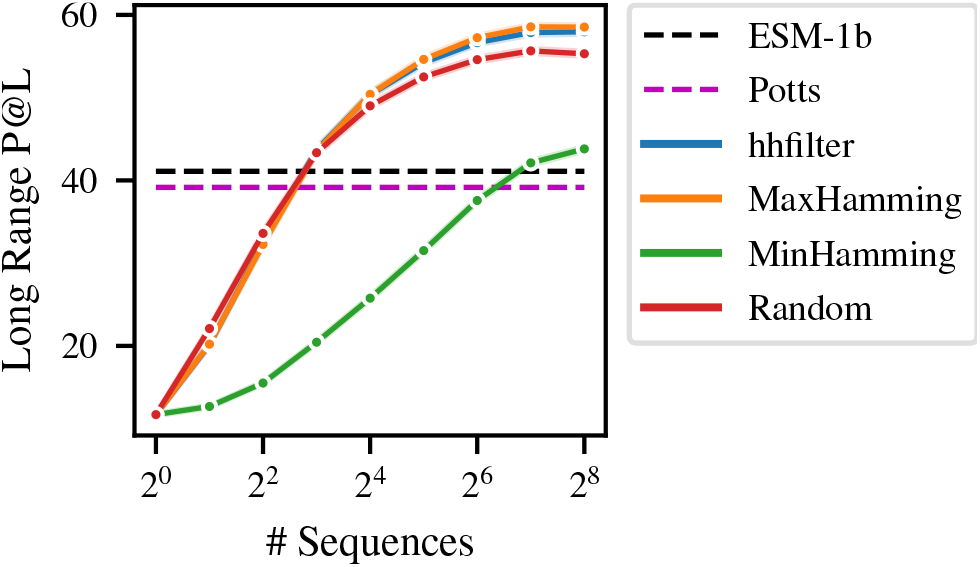
Comparison of MSA selection strategies. Model performance increases with more sequences. Selection strategies that maximize diversity of the input (MaxHamming and hhfilter) perform best. Random selection is nearly as good, suggesting the model has learned to compensate for the varying diversity during training time. Minimizing diversity performs worst. Using diversity maximizing approaches the MSA Transformer outperforms ESM-1b and Potts baselines using just 16 input sequences.

In principle, the model’s attention could allow it to identify and focus on the most informative parts of the input MSA. We find row attention heads that preferentially attend to highly variable columns. We also identify specific column attention heads that attend to more informative sequences. In this experiment random subsampling is used to select inputs for the model. Fig. 5 compares the distribution of attention weights with two measures of MSA diversity: (i) per-column entropy of the MSA; and (ii) computed sequence weights (Appendix A.5). Per column entropy gives a measure of how variable a position is in the MSA. Computed sequence weights measure how informative a sequence is in the context of the other sequences in the MSA. Sequences with few similar sequences receive high weights. The maximum average Pearson correlation between a row attention head and column entropy is 0.59. The maximum average Pearson correlation between a column attention head and sequence weights is 0.58. These correlations between attention weights and measures of MSA diversity suggest the model is specifically looking for informative sequences when processing the input.

**Figure 5.**
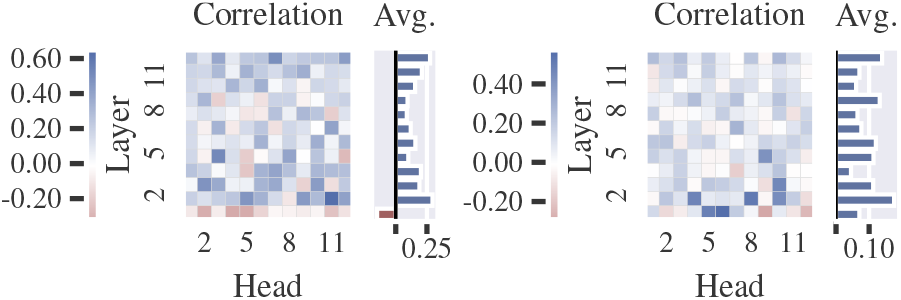
**Left:** Average correlation between row-attention and column entropy. This is computed by taking an average over the first dimension of each *L × L* row-attention map and computing correlation with per-column entropy of the MSA. **Right:** Average correlation between column-attention and sequence weights. This is computed by taking an average over the first two dimensions for each *L × M × M* column-attention map and computing correlation with sequence weights (see Appendix A.5). Both quantities are measures of MSA diversity. The relatively high correlation (> 0.57) of some attention heads to these measures suggests the model explicitly looks at diverse sequences.

**Figure 6.**
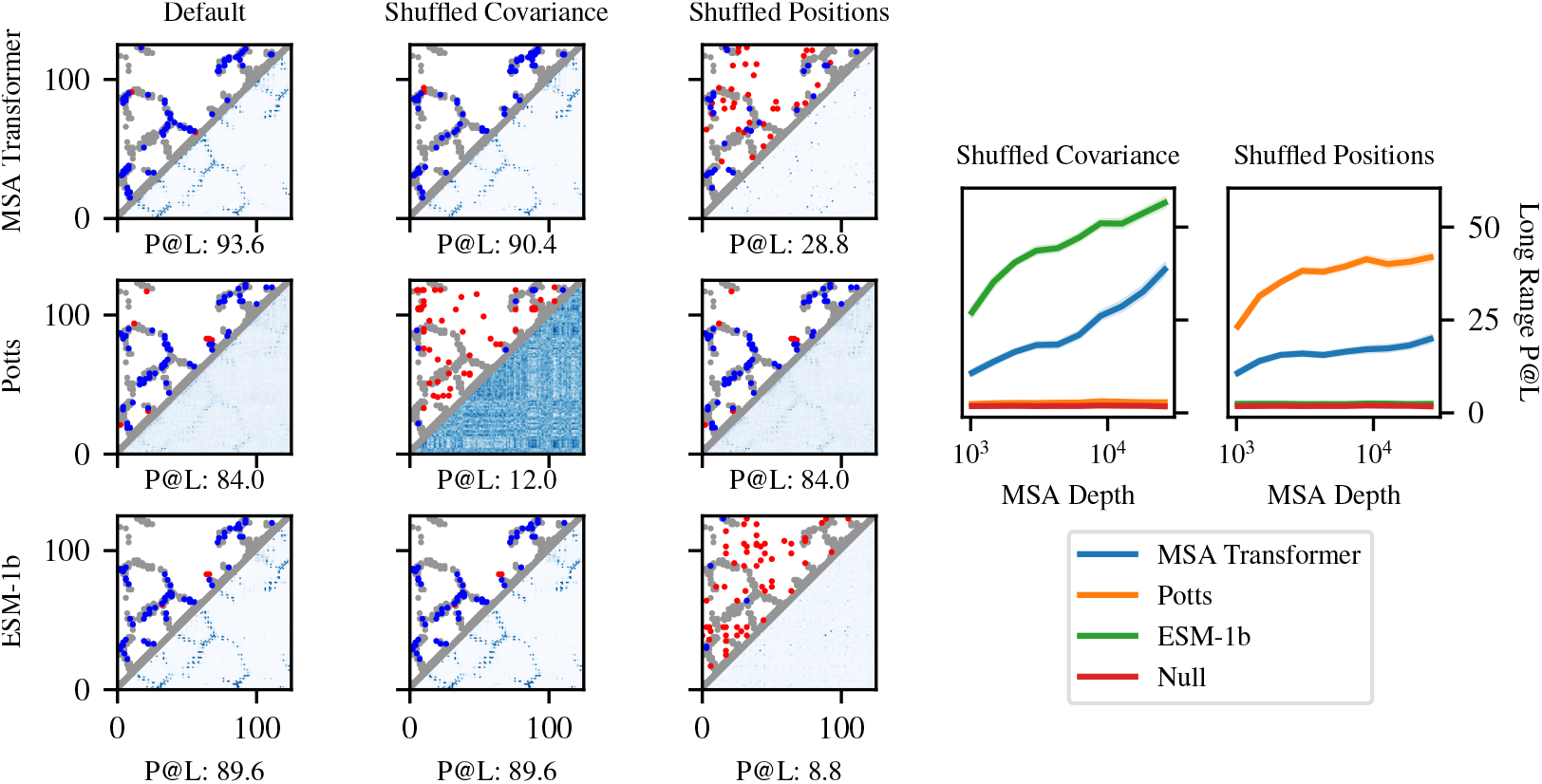
The MSA Transformer uses both covariance and similarity to training sequences to perform inference. **Left:** Examples (pdbid: 5ahw, chain: A) of model performance after independently shuffling each column of an MSA to destroy covariance information, and after independently permuting the order of positions to destroy sequence patterns. The MSA Transformer maintains reasonable performance under both conditions. A Potts model fails on the covariance-shuffled MSA, while a single-sequence language model (ESM-1b) fails on the position-shuffled sequence. **Right:** Model performance before and after shuffling, binned by depth of the original (non-subsampled) MSA. 1024 sequence selected with hhfilter are used as input to MSA Transformer and Potts models. MSAs with fewer than 1024 sequences are not considered in this analysis. Average Top-L long-range precision drops from 52.9 (no ablation) to 15.9 (shuffled covariance) and 27.9 (shuffled positions) respectively. A Null (random guessing) baseline is also considered. Potts model performance drops to the Null baseline under the first condition and ESM-1b performance drops to the Null baseline under the second condition. The MSA Transformer produces reasonable predictions under both scenarios, implying it uses both modes of inference.

### 5.2. Attention Corresponds to Protein Contacts

In Section 4.1, we use the heads in the model’s tied row attention directly to predict contacts in the protein’s three-dimensional folded structure. Following Rao et al. (2021) we fit a sparse logistic regression to the model’s row attention maps to identify heads that correspond with contacts. Fig. A.1 shows the weight values in the learned sparse logistic regression fit using 20 structures. A sparse subset (45 / 144) of heads are predictive of protein contacts. The most predictive heads are concentrated in the final layers.

### 5.3. Inference: Covariance vs. Sequence Patterns

Potts models and single-sequence language models predict protein contacts in fundamentally different ways. Potts models are trained on a single MSA; they extract information directly from the covariance between mutations in columns of the MSA. Single-sequence language models do not have access to the MSA, and instead make predictions based on patterns seen during training. The MSA Transformer may use both covariance-based and pattern-based inference. To disentangle the two modes, we independently ablate the covariance and sequence patterns in the model’s input via random shuffling. To ensure that there is enough information in the input for covariance-based extraction to succeed, we subsample each MSA to 1024 sequences using hhfilter, using only MSAs with at least 1024 sequences, and apply the model to unshuffled and shuffled inputs.

To remove covariance information, we randomly permute the values in each column of the MSA. This preserves per-column amino acid frequencies (PSSM information) while destroying pairwise correlations between columns. Under this condition, Potts model performance drops to the random guess baseline. Since ESM-1b takes a single sequence as input, the permutation trivially produces the same sequence, and the result is unaffected. Unlike the Potts model, the MSA Transformer retains some ability to predict contacts, which increases sharply as a function of MSA Depth. This indicates that the model can make predictions from patterns in the sequence profile in the absence of covariance.

To remove sequence patterns seen during training, we randomly permute the order of positions (columns) in the MSA. This preserves all covariance information between pairs of columns, but results in a scrambled input dissimilar to a real protein. Under this condition, Potts model performance is unaffected since its parameterization is invariant to sequence order. ESM-1b performance drops to the random guess baseline. The MSA Transformer does depend on sequence order, and predicts spurious contacts along the diagonal of the reordered sequence. When predicted contacts with sequence separation < 6 are removed, the remaining predictions align with the correct contacts. This shows the model can predict directly from covariance when presented with sequence patterns unobserved in training.

Together these ablations independently destroy the information used by Potts models and single-sequence language models, respectively. Under both conditions, the MSA Transformer maintains some capability to predict contacts, demonstrating that it uses both modes of inference.

## 6. Discussion

Prior work in unsupervised protein language modeling has focused on inference from individual sequences. We study an approach to perform inference from a set of aligned sequences in an MSA. We use axial attention to efficiently parameterize attention over the rows and columns of the MSA. This approach enables the model to extract information from dependencies in the input set and generalize patterns across MSAs. We find the internal representations of the model enable state-of-the-art unsupervised structure learning with an order of magnitude fewer parameters than current protein language models.

While supervised methods have produced breakthrough results for protein structure prediction (Jumper et al., 2020), unsupervised learning provides a way to extract the information contained in massive datasets of sequences produced by low cost gene sequencing. Unsupervised methods can learn from billions of sequences, enabling generalization to regions of sequence space not covered by structural knowledge.

Models fit to MSAs are widely used in computational biology including in applications such as fitness landscape prediction (Riesselman et al., 2018), pathogenicity prediction (Sundaram et al., 2018; Frazer et al., 2020), remote homology detection (Hou et al., 2018), and protein design (Russ et al., 2020). The improvements we observe for structure learning suggest the unsupervised language modeling approach here could also apply to these problems.

Improvement in unsupervised learning of structure and function with protein language models has been linked to scale of the models (Rives et al., 2020). Further scaling the approach studied here in the number of parameters and input sequences is a potential direction for investigating the limits of unsupervised learning for protein sequences.

## Acknowledgements

We thank Nicholas Bhattacharya, Zeming Lin, Sergey Ovchinnikov, and Neil Thomas for valuable input on the paper. Additionally, we acknowledge funding from Facebook, Berkeley Deep Drive, and DARPA XAI.

## A. Appendix

## A.1. Unsupervised Contact Prediction

For unsupervised contact prediction, we adopt the methodology from Rao et al. (2021), which shows that sparse logistic regression trained on the attention maps of a single-sequence transformer is sufficient to predict protein contacts using a small number (between 1 − 20) of training structures. To predict the probability of contact between amino acids at position *i* and *j*, the attention maps from each layer and head are independently symmetrized and corrected with APC (Dunn et al., 2008). The input features are then the values 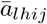 for each layer *l* and head *h*. The models have 12 layers and 12 heads for a total of 144 attention heads.

**Figure A.1.**
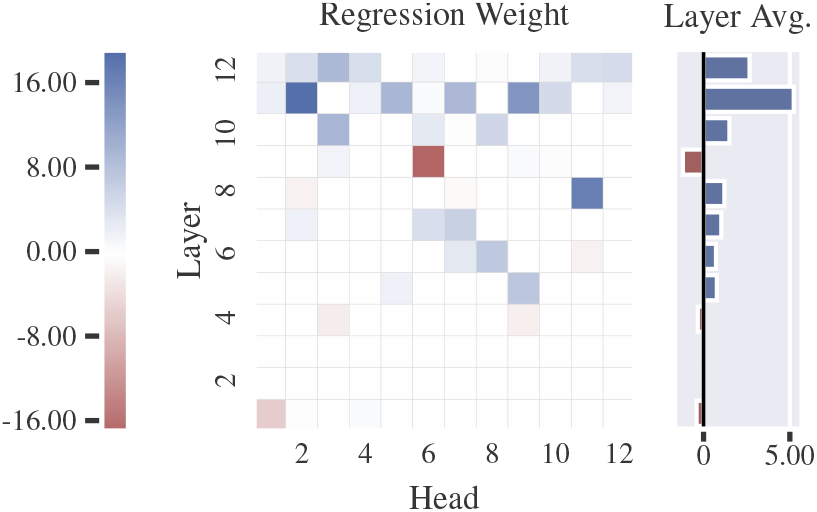
Weight values of learned sparse logistic regression trained on 20 structures. A sparse subset (55 / 144) of contact heads, largely in the final layers, are predictive of protein contacts.

An L1-regularization coefficient of 0.15 is applied. The regression is trained on all contacts with sequence separation ≥ 6. 20 structures are used for training. Trained regression weights are shown in Fig. A.1.

## A.2. Dataset Generation

For the unsupervised training set we retrieve the UniRef-50 (**?**) database dated 2018-03. The UniRef50 clusters are partitioned randomly in 90% train and 10% test sets. For each sequence, we construct an MSA using HHblits, version 3.1.0. (Steinegger et al., 2019) against the UniClust30_2017−10_ database (Mirdita et al., 2017). Default settings are used for HHblits except for the the number of search iterations (−n), which we set to 3.

## A.3. Ablation Studies

Ablation studies are conducted over a set of seven hyperparameters listed in Table. A.2. Since the cost of an exhaustive search over all combinations of hyperparameters is prohibitive, we instead train an exhaustive search over four of the hyperparameters (embedding size, block order, tied attention, and masking pattern) for 10k updates. The best run is then selected as the base hyperparameter setting for the full ablation study, in which only one parameter is changed at a time.

**Figure A.2.**
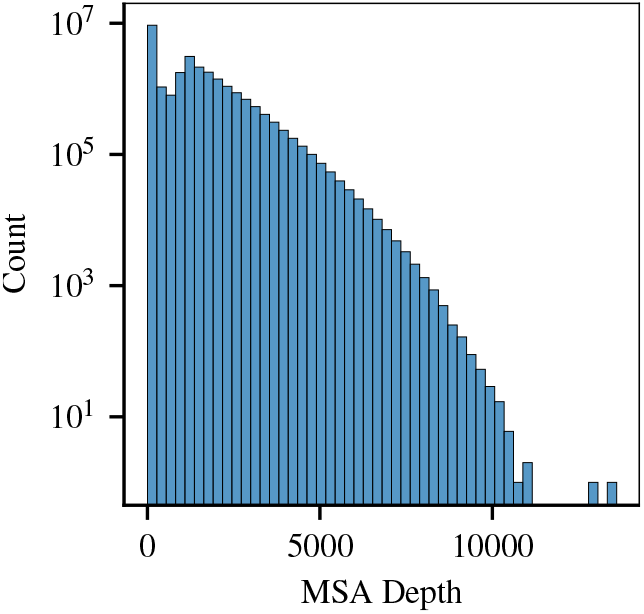
Distribution of MSA depths in the MSA Transformer training set. Average MSA depth is 1192 and median MSA depth is 1101.

**Table. A.1.** Validation perplexity and denoising accuracy on UniRef50 validation MSAs. PSSM probabilities and nearest-neighbor matching are used as baselines. To compute perplexity under the PSSM, we construct PSSMs using the input MSA, taking the cross-entropy between the PSSM and a one-hot encoding of the masked amino acid. When calculating PSSM probabilities, we search over pseudocounts in the range [10^−10^, 10), and select 10^−2^, which minimizes perplexity. For denoising accuracy, the argmax for each column is used. For nearest-neighbor matching, masked tokens are predicted using the values from the sequence with minimum hamming distance to the masked sequence. This does not provide a probability distribution, so perplexity cannot be calculated. MSAs with depth 1 are ignored, since the baselines fail in this condition. Perplexity ranges from 1 for a perfect model to 21 for a uniform model selecting over the common amino acids and gap token.

| Model | Perplexity | Denoising Accuracy |
| --- | --- | --- |
| PSSM | 14.1 | 41.4 |
| Nearest-Neighbor | - | 46.7 |
| MSA Transformer | **2.44** | **64.0** |

For the full ablation study, each model is trained for 100k updates using a batch size of 512. The four best performing models are then further trained to 150k updates. Contact prediction on the trRosetta dataset (Yang et al., 2019) is used as a validation task. Precision after 100k updates (and 150k for the best models) is reported in Table. A.2 and the full training curves are shown in Fig. A.3. The model with best hyperparameters is then further trained to 450k updates. The performance of this model is reported in Table. A.3. Validation perplexity is also reported in Table. A.2. In general we find limited correspondence between perplexity and contact prediction performance across models.

Potts (Balakrishnan et al., 2011), TAPE transformer (Rao et al., 2019), ESM-1b (Rives et al., 2020), ProtBERT-BFD, and ProTrans-T5 (Elnaggar et al., 2020) are used as unsupervised contact prediction comparisons. The best MSA Transformer outperforms all other methods by a wide margin, increasing long-range precision at L by a full 16 points. See below for a discussion of all seven hyperparameters.

## A.3.1. Embedding Size (*D*)

Since the MSA Transformer is provided with more information than single sequence protein language models, it is possible that many fewer parameters are needed to learn the data distribution. To test this hypothesis we train a model with half the embedding size (384 instead of 768) resulting in 30M total parameters. The resulting model achieves a Top-L long-range precision of 52.8 after 100k updates, 3.5 points lower than the baseline model. This suggests that model size is still an important factor in contact precision, although also shows that a model with fewer than 30M parameters can still outperform 650M and 3B parameter single-sequence models.

**Table A.2.**
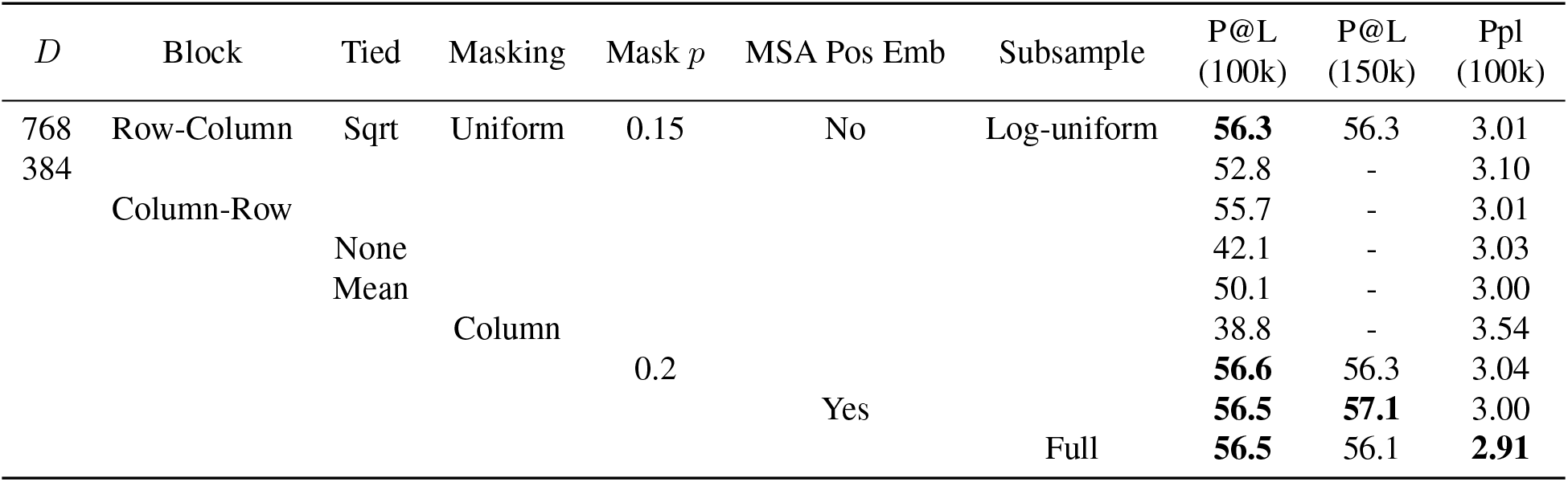
Hyperparameter search on MSA Transformer. P@L is long-range (*s* ≥ 24) precision on unsupervised contact prediction following Rao et al. (2021). Perplexity is reported after 100k updates and precision is reported after 100k and 150k updates.

**Table A.3.** Average precision on 14842 test structures for MSA and single-sequence models trained on 20 structures.

| Model | $6 \leq \text{sep} < 12$ | | | $12 \leq \text{sep} < 24$ | | | $24 \leq \text{sep}$ | | |
| --- | --- | --- | --- | --- | --- | --- | --- | --- | --- |
|  | L | L/2 | L/5 | L | L/2 | L/5 | L | L/2 | L/5 |
| Potts | 17.2 | 26.7 | 44.4 | 21.1 | 33.3 | 52.3 | 39.3 | 52.2 | 62.8 |
| TAPE | 9.9 | 12.3 | 16.4 | 10.0 | 12.6 | 16.6 | 11.2 | 14.0 | 17.9 |
| ProtBERT-BFD | 20.4 | 30.7 | 48.4 | 24.3 | 35.5 | 52.0 | 34.1 | 45.0 | 57.4 |
| ProTrans-T5 | 20.1 | 30.6 | 48.5 | 24.6 | 36.1 | 52.4 | 35.6 | 46.1 | 57.8 |
| ESM-1b | 21.6 | 33.2 | 52.7 | 26.2 | 38.6 | 56.4 | 41.1 | 53.3 | 66.1 |
| MSA Transformer | **25.6** | **41.0** | **64.6** | **31.9** | **48.9** | **71.1** | **57.4** | **71.7** | **82.1** |

**Table A.4.** Supervised Contact Prediction performance on CASP13-FM and CAMEO-hard targets. Reported numbers are long-range (*s* ≥ 24) contact precision. Three variants of the MSA Transformer are included for comparison: ^∗^unsupervised model, ^†^supervised model using final hidden representations of the reference sequence as input, ^‡^supervised model using final hidden representations of reference sequence and all attention maps as input. Baseline and final trRosetta models are also included for comparison. L is defined as the number of valid residues.

| Model | CASP13-FM |  |  | CAMEO |  |  |
| --- | --- | --- | --- | --- | --- | --- |
|  | L | L/2 | L/5 | L | L/2 | L/5 |
| Co-evolutionary | 40.1 | 52.5 | 65.2 | 47.3 | 60.9 | 72.1 |
| Unirep | 11.2 | 14.5 | 16.6 | 17.8 | 23.0 | 30.8 |
| SeqVec | 13.8 | 18.3 | 21.9 | 22.5 | 30.3 | 39.8 |
| TAPE | 12.3 | 14.4 | 17.8 | 15.9 | 20.6 | 26 |
| ProtBERT-BFD | 24.7 | 32.1 | 40.6 | 37.0 | 48.1 | 60.0 |
| ProTrans-T5 | 25.0 | 32.9 | 41.4 | 40.8 | 52.5 | 63.3 |
| ESM-1b | 28.2 | 37.4 | 50.2 | 44.4 | 57.2 | 68.4 |
| trRosetta<sub>base</sub> | 45.7 | 58.4 | 69.6 | 50.9 | 64.6 | 75.5 |
| trRosetta<sub>full</sub> | 51.8 | 66.6 | 80.1 | 53.2 | 67.1 | 77.5 |
| MSA Transformer* | 44.8 | 59.7 | 72.5 | 43.5 | 55.9 | 66.8 |
| MSA Transformer<sup>†</sup> | 54.5 | **70.0** | **80.2** | 53.6 | 68.4 | 78.0 |
| MSA Transformer<sup>‡</sup> | **54.6** | 68.4 | 77.5 | **55.8** | **69.8** | **79.1** |

## A.3.2. Masking Pattern

We consider two strategies for applying BERT masking to the MSA: uniform and column. Uniform masking applies masking uniformly at random across the MSA. Column masking always masks full columns of the MSA. This makes the training objective substantially more difficult since the model cannot look within a column of an MSA for information about masked tokens. We find that column masking is significantly worse (by almost 20 points) than uniform masking.

## A.3.3. Block Ordering

Row attention followed by column attention slightly outperforms column attention followed by row attention.

## A.3.4. Tied Attention

We consider three strategies for row attention: untied, mean normalization, and square root normalization (see Section 3). We find that tied attention substantially outperforms untied attention and that square root normalization outperforms mean normalization.

## A.3.5. Masking Percentage

As the MSA Transformer has more context than single sequence models, its training objective is substantially easier than that of single sequence models. Therefore, we explore whether increasing the masking percentage (and thereby increasing task difficulty) would improve the model. However, we do not find a statistically significant difference between masking 15% or 20% of the positions. Therefore, we use a masking percentage of 15% in all other studies for consistency with ESM-1b and previous masked language models.

**Figure A.3.**
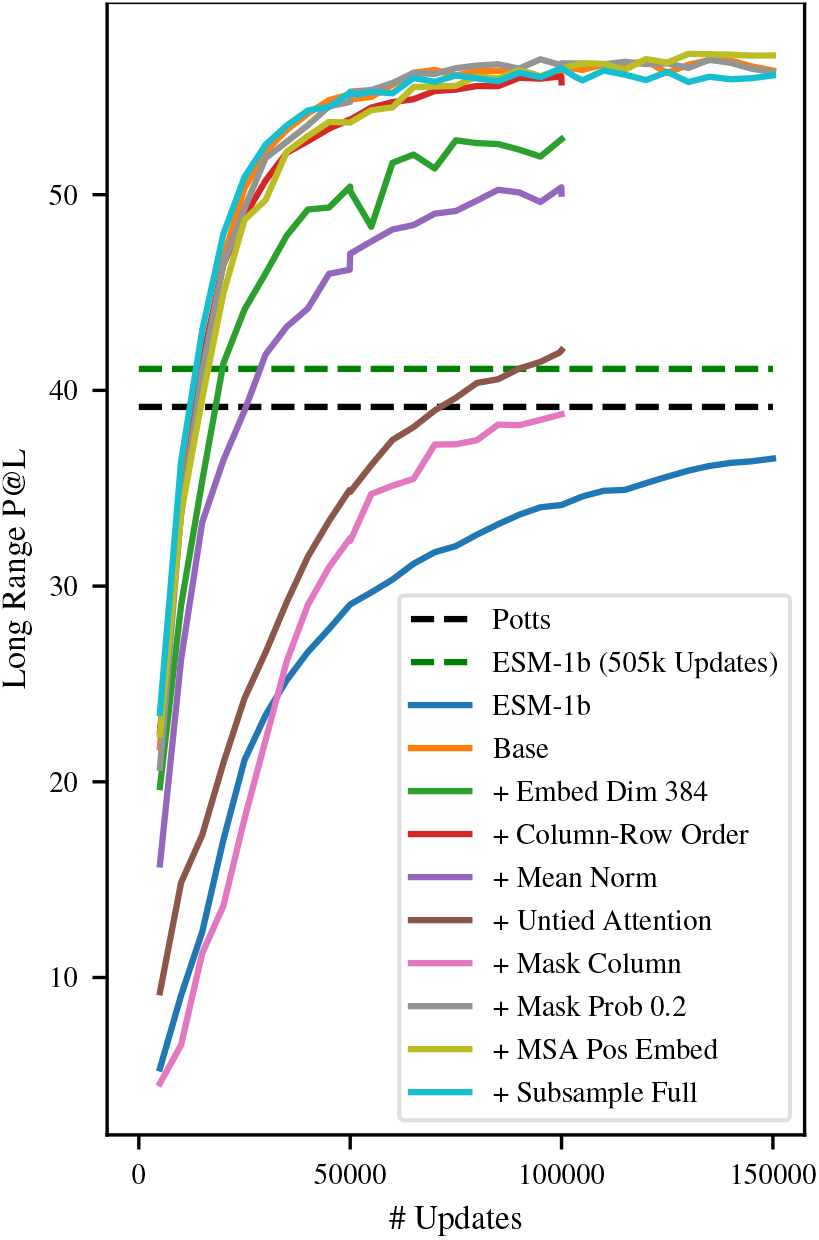
Training curves for MSA Transformer with different hyperparameters. See Section 4.4 for a description of each hyper-parameter searched over. ESM-1b training curve, ESM-1b final performance (after 505k updates), and average Potts performance are included as dashed lines for comparison.

## A.3.6. MSA Positional Embedding

An MSA is an unordered set of sequences. However, due to the tools used to construct MSAs, there may be some pattern to the ordering of sequences in the MSA. We therefore examine the use of a learned MSA positional embedding in addition to the existing learned sequence positional embedding. The positional embedding for a sequence is then a learned function of its position in the input MSA (not in the full MSA). Subsampled sequences in the input MSA are sorted according to their relative ordering in the full MSA. We find that the inclusion of an MSA positional embedding does modestly increase model performance, and therefore include it in our final model.

**Figure A.4.**
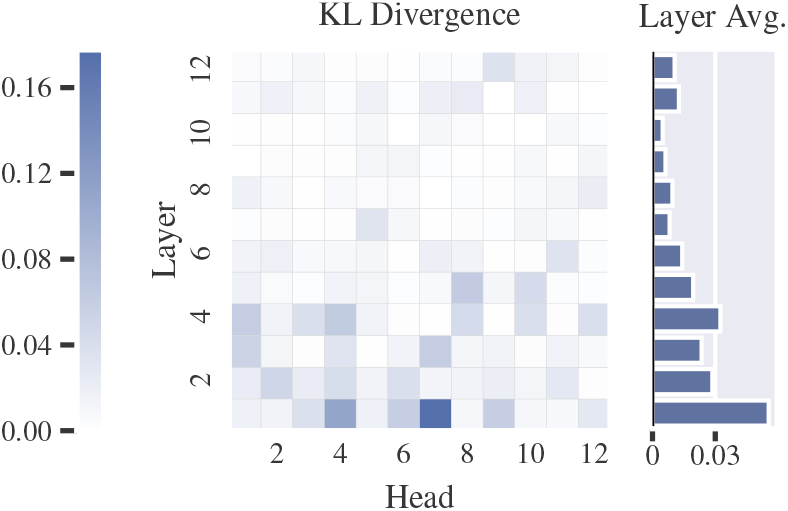
KL Divergence between distribution of row attention across amino acids and background distribution of amino acids. The fraction of attention on an amino acid *k* is defined as the average over the dataset of 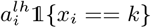, where *x*_*i*_ is a particular token in the input MSA and *a*^*lh*^ is the attention in a particular layer and head. KL Divergence is large for early layers but decreases in later layers.

## A.3.7. Subsample Strategy

At training time we explore two subsampling strategies. The first strategy is adapted from Yang et al. (2019): we sample the number of output sequences from a log-uniform distribution, with a maximum of *N/L* sequences to avoid exceeding the maximum tokens we are able to fit in GPU memory. Then, we sample that number of sequences uniformly from the MSA, ensuring that the reference sequence is always chosen. In the second strategy, we always sample the full *N/L* sequences from the MSA. In our hyperparameter search, most models use the first strategy, while our final model uses the second. We find no statistically significant difference in performance between the two strategies. However, it is possible that the log-uniform strategy would help prevent overfitting and ultimately perform better after more training.

The CCMpred implementation of Potts (Balakrishnan et al., 2011; Ekeberg et al., 2013), UniRep (Alley et al., 2019), SeqVec (Heinzinger et al., 2019), TAPE transformer (Rao et al., 2019), ESM-1b (Rives et al., 2020), ProtBERT-BFD, and ProTrans-T5 (Elnaggar et al., 2020) are used as supervised contact prediction comparisons. In Table. A.4 we show the complete results for long-range precision over the CASP-13 FM targets and CAMEO-hard domains referenced in (Yang et al., 2019). All baseline models are trained for 200 epochs with a batch size of 4.

## A.4. Attention to Amino Acids

**?** examine the distribution of amino acids attended to by single-sequence models. The attention in single-sequence models is roughly equivalent to the row-attention in our model, but there is no column-attention analogue. We therefore examine the distribution of amino acids attended to by the column attention heads. In Fig. A.4 we show the KL-divergence between the distribution of attention across amino acids and the background distribution of amino acids. The divergence is large for earlier layers in the model but decreases in later layers, suggesting the model stops focusing on the amino acid identities in favor of focusing on other properties.

## A.5. Sequence Weights

Sequence reweighting is a common technique used for fitting Potts models which helps to compensate for data bias in MSAs (Morcos et al., 2011). Informally, sequence reweighting downweights groups of highly similar sequences to prevent them from having as large of an effect on the model. The sequence weight *w*_*i*_ is defined as,

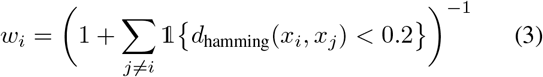

where *x*_*i*_, *x*_*j*_ are the *i*-th and *j*-th sequences in the MSA, *d*_hamming_ is the hamming distance between two sequences normalized by sequence length, and *w*_*i*_ is the sequence weight of the *i*-th sequence.

## Footnotes

1 The final vocab size is 29, consisting of 20 standard amino acids, 5 non-standard amino acids, the alignment character ‘.’, gap character ‘-’, the start token, and the mask token

